# OLIVER: A Tool for Visual Data Analysis on Longitudinal Plant Phenomics Data

**DOI:** 10.1101/411595

**Authors:** Oliver L Tessmer, David M Kramer, Jin Chen

**Affiliations:** Dept. of Energy Plant Research Lab, Michigan State University, East Lansing, USA; Inst. for Biomedical Informatics, University of Kentucky, Lexington, USA

**Keywords:** phenomics, instant data visualization, heat map, hypothesis statistical test

## Abstract

There is a critical unmet need for new tools to analyze and understand “big data” in the biological sciences where breakthroughs come from connecting massive genomics data with complex phenomics data. By integrating instant data visualization and statistical hypothesis testing, we have developed a new tool called OLIVER for phenomics visual data analysis with a unique function that any user adjustment will trigger real-time display updates for any affected elements in the workspace. By visualizing and analyzing omics data with OLIVER, biomedical researchers can quickly generate hypotheses and then test their thoughts within the same tool, leading to efficient knowledge discovery from complex, multi-dimensional biological data. The practice of OLIVER on multiple plant phenotyping experiments has shown that OLIVER can facilitate scientific discoveries. In the use case of OLIVER for large-scale plant phenotyping, a quick visualization identified emergent phenotypes that are highly transient and heterogeneous. The unique circular heat map with false-color plant images also indicates that such emergent phenotypes appear in different leaves under different conditions, suggesting that such previously unseen processes are critical for plant responses to dynamic environments.

## I. INTRODUCTION

Modern genetic tools have revolutionized biology [28], [33]. It is now possible to rapidly and inexpensively sequence the genomes of any organism on the planet [29], [34]. However, this huge dataset does not, by itself, answer most major questions in biology, because we do not know how these genes control complex biological processes. The next revolution in biology will be to connect the genomes to the phenomes of organisms, i.e. to understand how the genome and its related metagenome control complex behaviors [4], [46]. To achieve this integration, we need to measure and interpret multiple phenotypes (i.e., the specific properties and behaviors of organisms) under a range of dynamic environmental conditions (or stimuli) in a large number of genetic variants [14], [23]. This is a hyper-complex problem requiring new approaches to measurements, data analyses and the generation of functional insights. Recent developments in plant photosynthesis phenotyping make it possible to probe plant growth, photosynthesis and other properties under dynamic environmental conditions [7], [11], [18], [31], [44]. Similar approaches are impacting other fields, such as biochemistry, drug development and behavior studies [3], [5], [6], [40].

Despite these major phenotyping technological advances, biomedical scientists are facing the difficulty of analyzing longitudinal phenomics data, for the nonlinear temporal patterns in a high-dimensional space are difficult to detect. Particularly, plant photosynthetic phenomics experiments are often conducted under dynamic environmental conditions, in order to measure the influence of environment (e.g. abrupt changes in light intensity or temperature) on photosynthesis (e.g. excessive losses in photosynthetic efficiency and/or increases in photodamage). In plants, photosynthesis must respond to changing environment to provide the optimal amount of energy to meet the needs of the organism, in the correct forms, without producing toxic byproducts, e.g. reactive oxygen species or glycolate [13], [24], [47], [50]. In this context, photosynthesis can be viewed as a set of integrated modules that form a self-regulating network that is sensitive to changes in both environmental parameters (e.g. light intensity) and metabolic or physiological factors [22]. Consequently, it is crucial to *simultaneously* model genotypes, phenotypes, and environmental factors and identify complex relationships among multiple variables to arrive at a holistic characterization of plant performance [50].

In practice, for plant phenomics data analysis, a combination of visual and statistical data analytics approach emerges [7]. The concept of visual knowledge discovery, which has been extensively utilized in a wide variety of data mining and machine learning applications [10], [15]–[17], [42], adopts the unique human perceptual and cognitive abilities and combines it with modern machine learning techniques, leading to accelerated multi-dimensional pattern recognition [21].

A critical component of visual and statistical data analysis is visual data representation. Due to the limitation of human cognition and perception, the quantity of information a person can examine at a moment is limited, which significantly affects the volume of data a user can view and consequently the pattern a person can retrieve for any specific purposes [45], [48]. A *clean, simple and interactive phenomics data visualization interface* is required for understanding, comparing, and analyzing complicated relationships among individuals in a multi-dimensional space [8], [20]. With the phenomics data visualization interface, users can effectively gaining insights from big data to interpret complex behaviors, to determine which genes are responsible to certain phenotypes under specific environmental conditions, or to identify genes involved in certain biological process and test mechanistic hypotheses.

To facilitate visual phenomics knowledge discovery, we present an interactive phenomics data visualization tool called **OLIVER**, which is the acronym of five key processes in visual data analysis, i.e. Observe, Link, Integrate, Verify, Explore, and Reveal (see the main interface in Figure 1 and the workflow in Figure 2.). OLIVER visually represents data using interactive animated heat maps. Comparing with the existing data visualization tools such as KaleidaGraph [43], Origin, and Microsoft Excel that are often used to generate heat maps, OLIVER is advantageous on its design principles, i.e. *any user adjustment will trigger real-time display updates for any affected elements in the workspace*. Note that not all the functions in OLIVER are conceptually new. In fact, functions like sorting or searching are available in almost every heat map tool, and amplification and color curve adjustment are common in many photo-editing tools such as Adobe Photoshop and GIMP. But OLIVER is the first tool to bring all these into one platform for visual data analysis. Furthermore, to our knowledge, the key of the heat map visualization is not to appropriately display a heat map, it is instead to *instantly display a heat map when adjustments (such as sorting, clustering, merging, splitting, re-coloring) are applied*. This enables the trial-and-error (or “what if?”) strategy in the data analysis workflow. It effectively increases the incidence of new discoveries (also called exploratory research or even serendipity) [36]. For example, the user may notice variations in a region that was assumed to be homogeneous as they click-and-drag the color scheme editor, leading to new discoveries on developmental regulation [8].

**Fig. 1.**
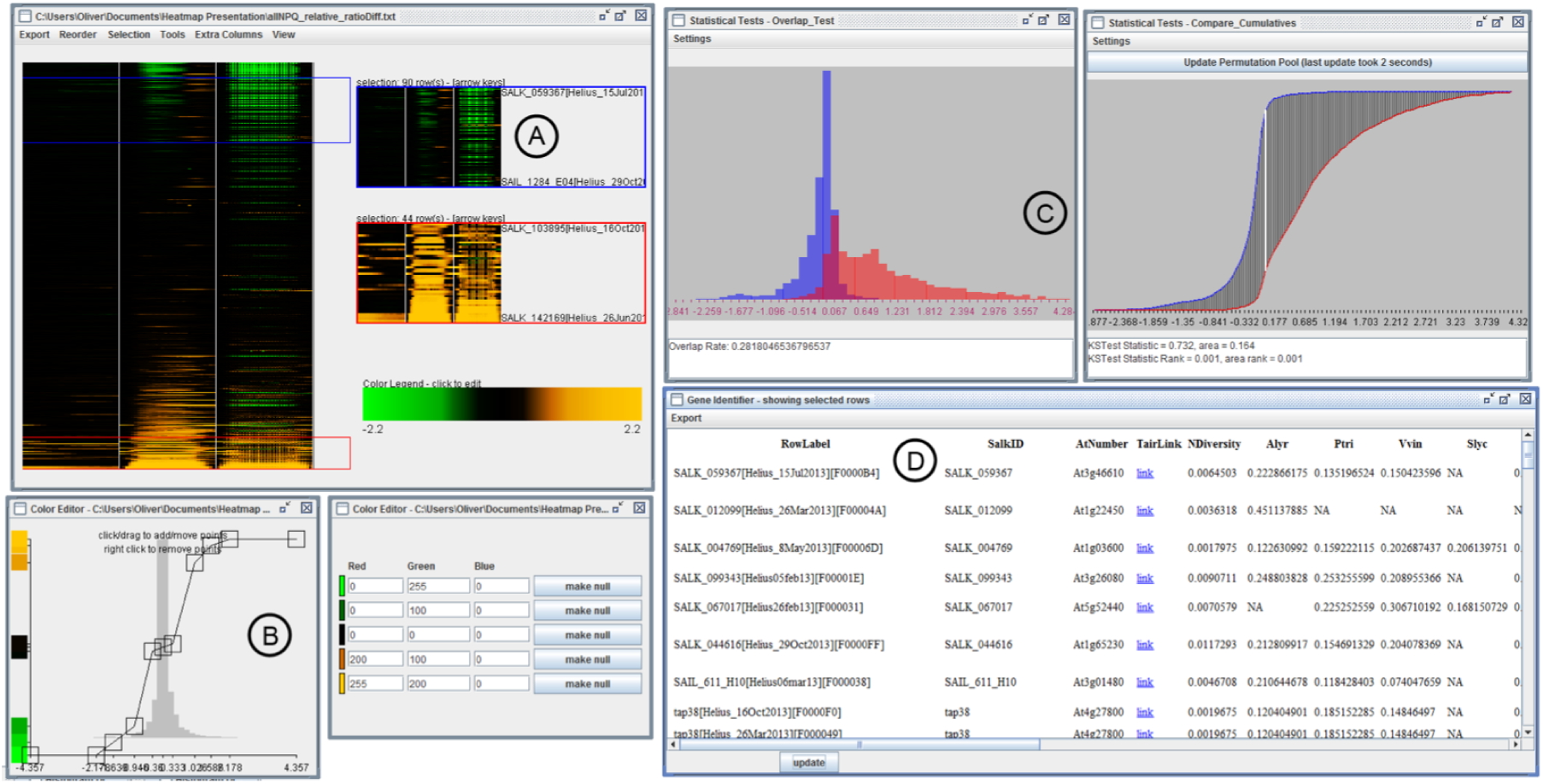
OLIVER workspace with (a) a phenotype-over-time heat map loaded from a spreadsheet, with two subsets of rows selected, (b) two dialogs for editing the color curve of the heat map, one to edit the RGB channel intensities (right), and the other to change the way the base colors are interpolated (left), (c) statistics dialogs showing overlaid histograms representing the frequency distributions of values in the selected subsets (left), as well as a comparison of their cumulative curves and a p-value (right), (d) the Gene Identifier dialog, which presents external data corresponding to a row selection in the heat map.

**Fig. 2.**
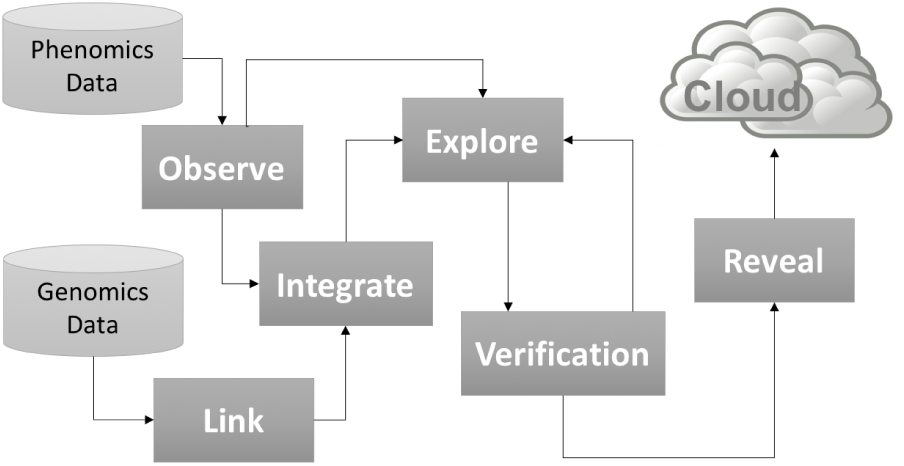
The workflow of OLIVER. OLIVER is the acronym of five key processes in visual data analysis: Observe, Link, Integrate, Verify, Explore, and Reveal.

**Fig. 3.**
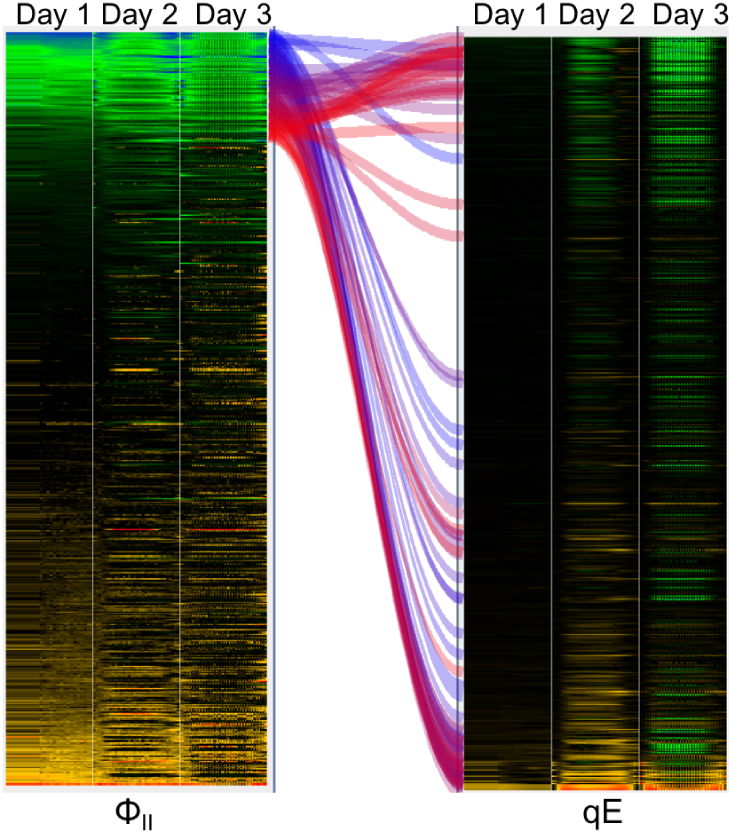
For a 3-day plant phenotyping experiment, two heat maps of photosynthetic phenotype parameters (i.e. photosynthetic rate of photosystems II Φ_*II*_ and the pH- or energy-dependent component of non-photochemical quenching *qE*) have their rows linked based on the matching row labels. A row-based selection is made based on the row order of the left heat map, and bands are automatically drawn between the selected rows and the corresponding rows in the other heat map.

In summary, OLIVER is a visual data analysis tool specially designed for big longitudinal phenomics data with a unique feature that any user adjustment will trigger real-time display updates, allowing biologists to quickly integrate and compare phenomics data resulting from large-scale phenotyping experiments, and genomics information in public databases.

## II. BACKGROUND

In the last decade, with the rapid development of data-driven techniques, many data visualization tools have been developed to visually recognize patterns and trends in big data and to facilitate statistical analysis and data mining [16], [35], [42]. Among all the data visualization techniques, a particularly useful tool is the heat map, which is a graphical representation of data making full use of color to extend perception into multiple dimensions by allowing properties to be mapped both spatially and as color gradients [49]. Specifically, the 2D heat map visualization allows researchers to quickly grasp the state and impact of a large number of variables at one time [37]. 2D heat map tools have been widely used in visualizing data in gridded formats, such as genome-wide gene expression or genetic mutations [30], [32].

While there are many available heat map programs for genomics data visualization [1], [2], [9], [12], [19], [39], to our knowledge, tools focused on large-scale phenomics data visualization is still absent. Phenomics data, in contrast to genomics data, are collected from numerous sources, represent vastly different phenomena, and are time and condition dependent [23]. In general, the rows and columns of a phenomics dataset are the individuals to test and the time points the individuals were phenotyped, respectively. Phenotype values change dynamically with environments or stimuli whereas the genome is constant, making phenotyping data highly complex and condition-dependent [7]. The relationships among testing objects (such as mutant lines), environmental conditions (such as light intensity and temperature), and the behavior of organisms are mutually interacting so that effects may be seen only under certain sets of conditions and genetic backgrounds [27].

In this article, we introduce OLIVER, which is designed to visualize and analyze complex phenotypic datasets in ways that reveal emerging phenomena and trends and thus to generate and test biological hypotheses and communicate results visually. OLIVER was developed with the general purpose of extracting knowledge from large-scale phenomics data. By developing the software in intimate collaboration between experts in biology, bioinformatics, and computer science, we have demonstrated that OLIVER is able to identify the insights of plant photosynthetic phenomics data [7].

## III. ARCHITECTURE

By integrating instant data visualization and hypothesis testing, we have developed a phenomics visual data analysis tool called **OLIVER**. OLIVER is the acronym of “Observe, Link, Integrate, Verify, Explore, and Reveal”, which are the six critical processes of visual knowledge discovery. The workflow of OLIVER is shown in Figure 2.

### A. OBSERVE Emerging Phenomena in Phenomics Data

The 2D heat map, the main view of OLIVER, is designed to instantly visualize large-scale time-resolved phenomics data and to assistant users to optimize data visualization. The core display functions are aimed towards presenting one or more entire phenomics data sets in the same workspace, providing essential operations for both broad and focused analysis.

In very long or large-scale phenomics experiments, such as to measure the growth rates of plants in their whole life span or to phenotype thousands of plants at the same time, the number of columns/rows could be even higher than the number of pixels in the width/height of the heat map display. To render large heat maps at real time, loss-less heat map snapshots are periodically generated in the memory. Computer graphics card(s) are used for rapid scaling/stretching these snapshots to any size to facilitate heat map visualization. When adjustments on the colors or the order of rows and columns are made by the users, real-time previews can be generated by sampling only the heat map cells that are visible on the screen.

In the main view, rows of a heat map can be sorted or clustered according to the columns that users manually or automatically select. In the sorting dialog, the unsorted and sorted heat maps are displayed side-by-side for comparison, and the sorted heat map will update in real-time as users click-and-drag to select the columns to sort (see Figure 6). A heat map can be instantly re-colored using an interactive color curve interface (similar to the curve dialog in Photoshop). In a heat map, appropriate coloring allows users to better sort or cluster the rows and the columns and to maximally differentiate phenotype values. With OLIVER’s interactive value-to-color mapping interface (see Figure 5), new color schemes can be rapidly developed and saved for future use. The purpose of re-coloring is to allow users to precisely control the value-to-color mapping, which is particularly useful to capture extreme values in non-normal distributions. In the re-coloring interface, color adjustments can be triggered using intuitive click-and-drag gestures. As one of the central features of OLIVER, any adjustment will trigger real-time display updates for any affected elements in the workspace.

**Fig. 4.**
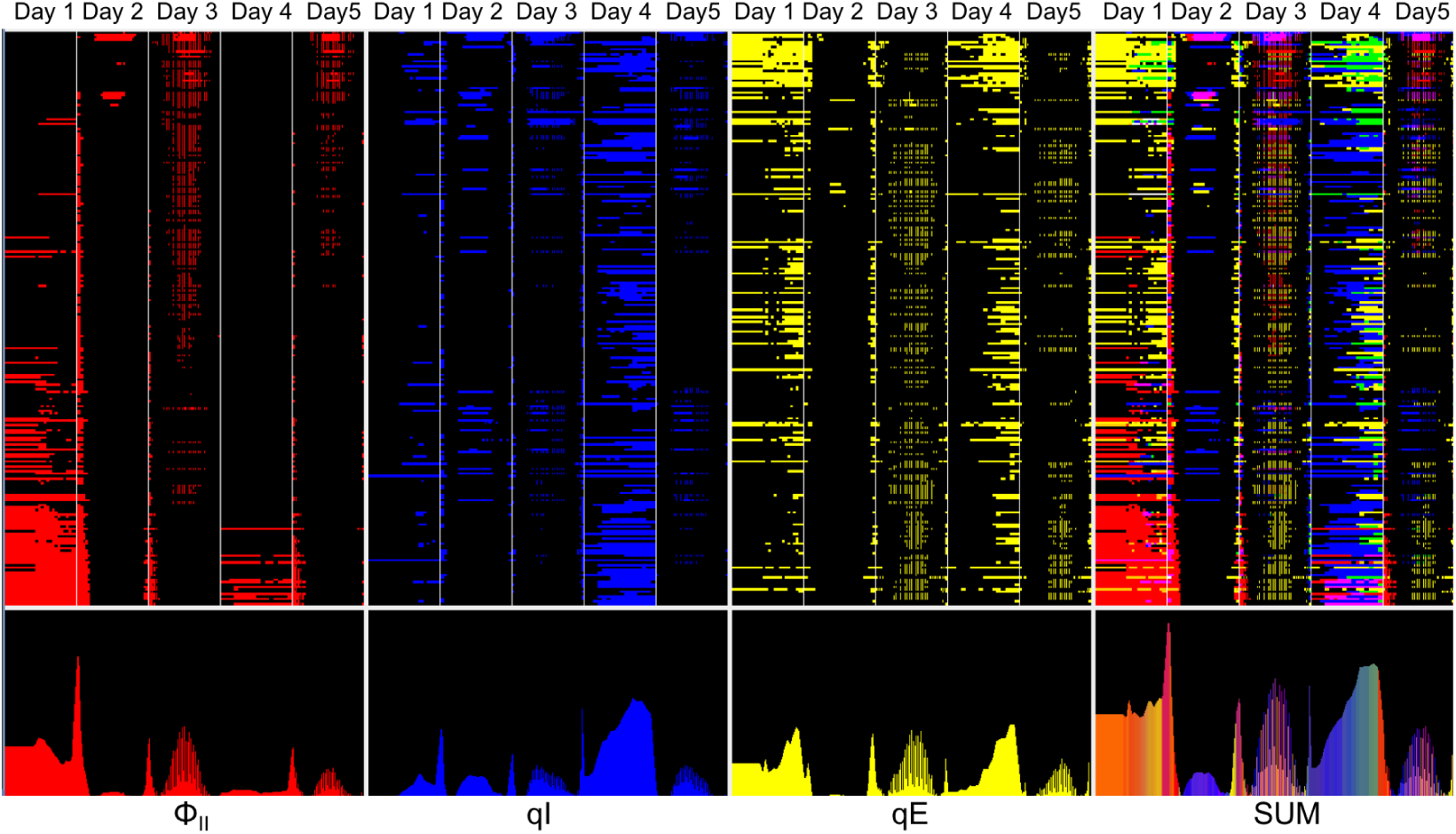
For a 5-day plant phenotyping experiment, the binarized heat maps on the left represent three different photosynthetic phenotype parameters (Φ_*II*_, *qI*, and *qE*). They were created by applying thresholds to the original heat maps. The heat map on the right is a combined visualization created by assigning logical colors to every possible combination of the three corresponding values on the left. On the bottom row, totals have been accumulated for each column to give a summarized view of expression-over-time for each phenotype parameter, as well as for the combined result. Users can adjust the binarization thresholds and instantly view the changes in this display.

**Fig. 5.**
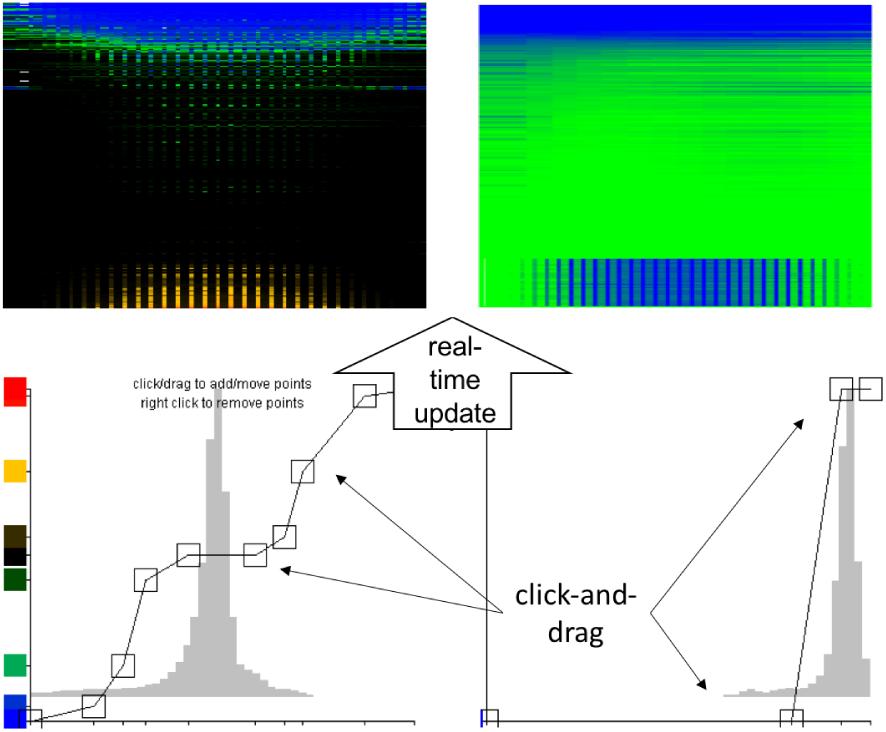
Color curve interface (bottom row) and the associated heat maps (top row). The right side features data with a single-tailed distribution, while the left side contains two tails. This interface features both manual and automated options for assigning an appropriate color scheme to a heat map.

**Fig. 6.**
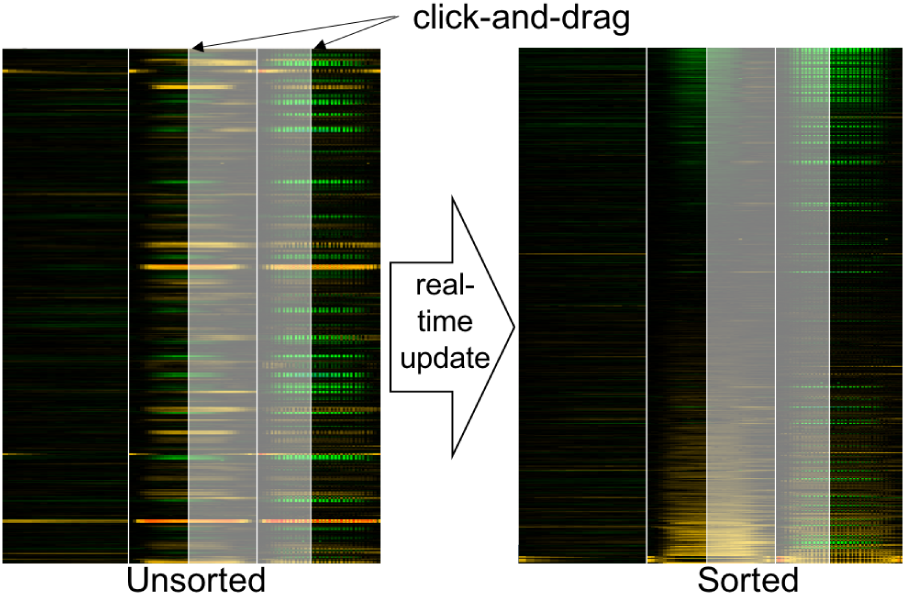
Sorting Dialog. Both the unsorted (left) and sorted (right) heat maps are displayed, where the heat map on the right updates in real-time as users click-and-drag to select the time-range to sort (highlighted in white).

Faithfully visualizing rows and columns in a heat map is imperative to gaining appropriate insights from sophisticated phenomics data. A unique property of phenotyping experiments is that, to obtain the data with the best quality, the phenotyping sampling rate often varies within the same experiment. Accordingly, the width of each column in the rendered heat map of OLIVER is set to be proportional to the phenotyping sampling rate. The OLIVER workspace can also display multiple heat maps simultaneously, allowing them to be independently rendered, ordered, and related to each other via common features. OLIVER also allows users to visualize heat maps at multiple levels of data granularity ranging from the averaged phenotypes of every cluster to the sample groups measured using different devices, to the view of individuals without losing any details, thus facilitating investigations at different biological aspects (see Figure 1a, red and blue boxes).

### B. LINK Metadata from External Sources

OLIVER can connect the input phenomics data with a range of online databases via external links to facilitate a better understanding of the phenomics data. As shown in Figure 1d fourth column, external links to the corresponding genes/proteins in the TAIR database [25] provide the most recently updated gene structure of Arabidopsis thaliana along with gene product, gene expression, DNA and seed stocks, genome maps, genetic and physical markers, and publications, which is particularly useful for Arabidopsis phenotyping experiments. Besides TAIR, the OLIVER API allows for integration with a variety of existing databases.

The other columns in Figure 1d show results generated based on the lookup-tables customized by users. Inspired by the modern object-relational database model [41], all the look-up tables in OLIVER are designed to be accessed in a chained fashion, i.e. the result of the look-up table loaded first becomes the input of the second one and so on.

The external data (such as sub-cellular localization in Gene Ontology), once being linked, can be represented visually as extra columns to a heat map (see Figure 8). Extra columns are typically rendered independent to the original heat map, and are given a unique color scheme to represent linked results that are either qualitative (e.g. the experimenter who generated the data for each row), or quantitative (e.g. the evolutionary rate of a protein for each row). Extra data can be saved in the same data file containing the input phenomics data for easier data management.

**Fig. 7.**
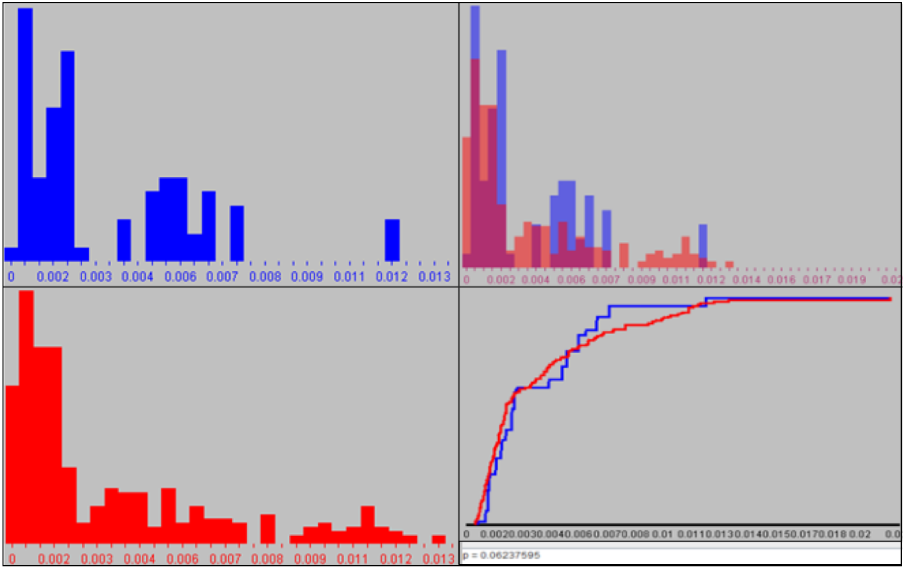
Color-coded frequency distributions of two heat maps (top and bottom-left), which are overlaid (top-right) and are compared using Kolmogorov-Smirnov test (bottom-right).

**Fig. 8.**
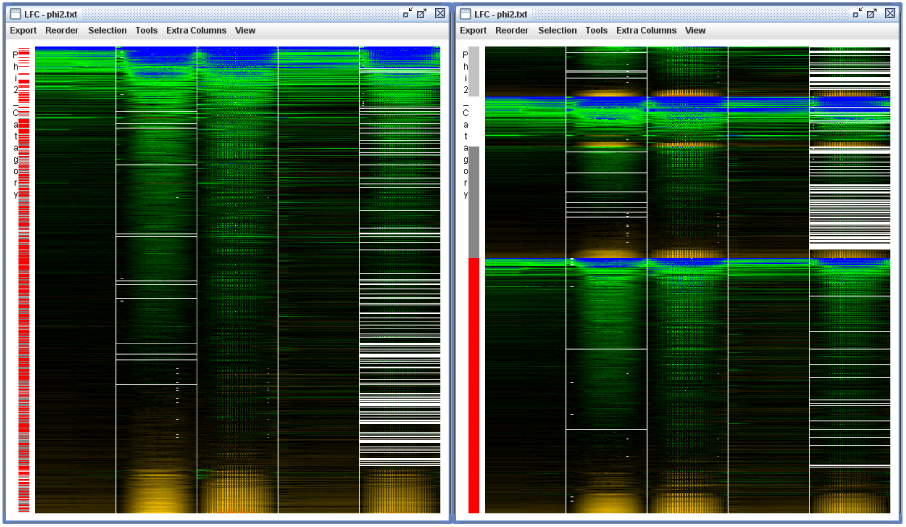
A heat map is shown on the left with rows ordered according to the averaged values, and one additional column containing categorical information pertaining to the individual row entries. On the right, the same heat map has been reordered such that the extra column categories take precedence in sorting. This allows users to explore the relationships between phenotype measurements and external information.

### C. INTEGRATE Results for Hypothesis Generation

The power of phenomics data analysis comes from the integration of multiple kinds of phenotype parameters (such as the photosynthetic rate of photosystems II Φ_*II*_ and the pH- or energy-dependent component of non-photochemical quenching *qE*, see Figure 3). This is because while a single phenotyping parameter describes a general effect, checking multiple phenotype parameters will lead to the identification of specific pathways and genes that are key to phenotype regulation. Thus phenomics data is inherently many-dimensional and difficult for humans to perceive.

OLIVER allows users to statistically test interactions among multiple heat maps, revealing trends in multi-dimensional phenomics datasets. In OLIVER, the data for each phenotype parameter can be visualized as a binary heat map (see Figure 4, left three heat maps). Then, the multiple binary heat maps can be combined into a single one, which is colored by assigning logical colors to every possible combinations of the corresponding values in binary heat maps, as shown in Figure 4, the right heat map. In the integration process, any operation (e.g. changing the percentages to mitigate the rate of loss-of-detail caused by binarization) triggers real-time display updates, allowing for rapid exploration of hypotheses considering multiple phenotype parameter combinations.

The integration function enables the discovery of new functional relationships between phenotype parameters. Figure 3 demonstrates one such case using an intuitive visualization. In the figure, a set of individuals with a single extreme phenotype value under one phenotype parameter (left) are linked to a second phenotype parameter (right), where the corresponding individuals are clustered on two different extremes.

### D. Rapid Statistical VERIFICATION of Hypotheses

By conducting on-the-fly statistical tests on phenomics data sets, OLIVER allows users to verify whether a hypothesis is statistically sound or not directly on the visualized results. To provide statistic tests, OLIVER takes advantage of R and its libraries in statistics. For example, OLIVER performs the Kolmogorov-Smirnov test [26] to check whether two input data have significantly different distributions. The result shown in Figure 7 (bottom-right) was generated by calling R function *ks.test* in the background [38]. In OLIVER, any modification that would affect the statistical tests (e.g. by changing the row or column of selection) will result in instant update in the statistical test.

### E. Visually EXPLORE Multiple Hypotheses

The seamless integration of all the modules in OLIVER allows users to test and compare multiple “what if” hypotheses. For instance, users can explore phenotype relationships by adjusting the color scheme of a heat map (Figure 1b) to highlight only values far from the control, and then reorder the rows (Figure 6) based on each row’s variance in a given temporal range. The resulting heat map may identify mutant lines with diverse responses over the course of a specific stimuli, leading to insights about genes that affect the given phenotypes. Furthermore, by taking advantage of heuristic options (such as preset color maps, automatic sorting, and scripting), this data processing workflow can be completed within a few minutes. This greatly increases the rate at which researchers can test their hypotheses.

OLIVER provides enough flexibility towards the design of workflow so that users are free to explore data at their own pace. OLIVER has a variety of functions allowing users to gain new insights by exploring the results of every step, or it can run all the steps and show the final results. Fast-paced users will reach end-goal visualizations as quickly as possible, while exploitative users will be provided with fully interactive in-depth visualizations for each step in an open-ended workflow.

### F. REVEAL New Knowledge by Sharing Complex Results

OLIVER provides a web-based module powered by Java applet to distribute its workspace in case-specific configurations, so that multiple users can collaboratively view and explore the same phenomics data. For instance, a user can create and share a workspace of OLIVER with a group of viewers. Then the viewers would be presented with a workspace populated with the same data using the same settings. In addition, the results of OLIVER can be saved in graphical forms that are readily interpretable by users, as well as files in the form of publishable graphics, spreadsheets, or the internal binary file types that OLIVER can reload. All of these files can be easily shared among users.

### G. Software Implementation

OLIVER was developed using Netbeans IDE 8.0, and compiled using JDK 1.8. Java Swing API was used exclusively for all display functions. OLIVER uses R and its statistics libraries [38] to provide a wide variety of statistical analysis. For plant phenotyping, OLIVER includes look-up tables that can link Arabidopsis mutant lines to the corresponding genes in the TAIR web site [25]. Here we introduce the implementation details of three key functions in OLIVER.

#### 1) Sorting and Clustering

Sorting and clustering are basic functions in OLIVER that provide users accessible options for re-ordering and grouping rows of a heat map. Users can sort a heat map based on phenotype value sums, standard deviations, absolute values, *etc*. Users can also group the rows using hierarchical or k-means clustering. If a look-up table is provided, sorting/clustering can be performed on the extra information in the look-up table in addition to the phenotype values in the heat map. A unique feature is that users can sort/cluster multiple linked heat maps synchronously, and conduct statistical tests on them as if they work on a single heat map. This allows users to explore the relationships among multiple phenotype parameters. Users can also propagate the row ordering of a heat map to other heat maps. Another unique feature is that users can preview sorting/clustering results before actually running the function in the entire data set (see Figure 6). A preview is generated and updated in real-time as the column/row selection is tweaked and the sorting/clustering methods is chosen. Users can instantly see the individuals being sorted or grouped and the results of statistical tests when they specify sorting/clustering/testing parameters.

#### 2) Zoomed View

While an input phenomics data set can be displayed on a global level, the nature of phenomics data analysis demands visualizing multiple levels of granularity simultaneously. To this end, an interactive “zoomed view” function has been implemented. Consider a phenomics data set containing multiple biological replicates for each genotype. The input data can be consolidated to an *averaged heat map*, where each row contains the averaged phenotype values for each genotype in the input. A row selection can be made conveniently by clicking-and-dragging the mouse over the heat map. The selection will instantly result in a zoomed view of the values in the selected area at a finer level of granularity. With the more detailed information provided by OLIVER, users can visually check sample variance, experiment consistency, and distribution of raw data (see Figure 9). Phenotype data are often extracted from images, and a measured phenotype value is often the average of many pixels. In this case, OLIVER supports yet another level of granularity, where a detailed image is presented for each cell in the heat map. For grey scale images, the same value-to-color mapping is adopted to create a false-color image (see Figure 11). This level of detail is helpful in understanding unexpected results or identifying errors in data collection.

**Fig. 9.**
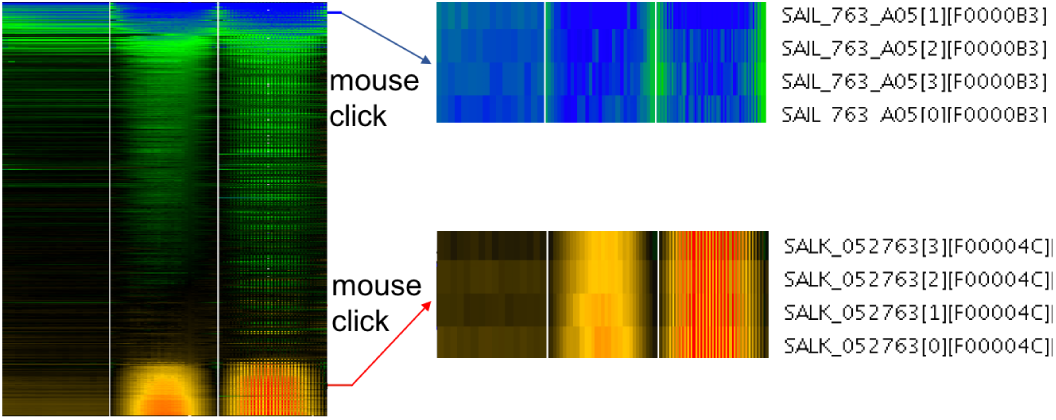
A phenotype-over-time heat map, where the values in the heat map are the averaged values of multiple biological replicates of each genotype. The right heat maps show the raw data of a genotype before averaging.

**Fig. 10.**
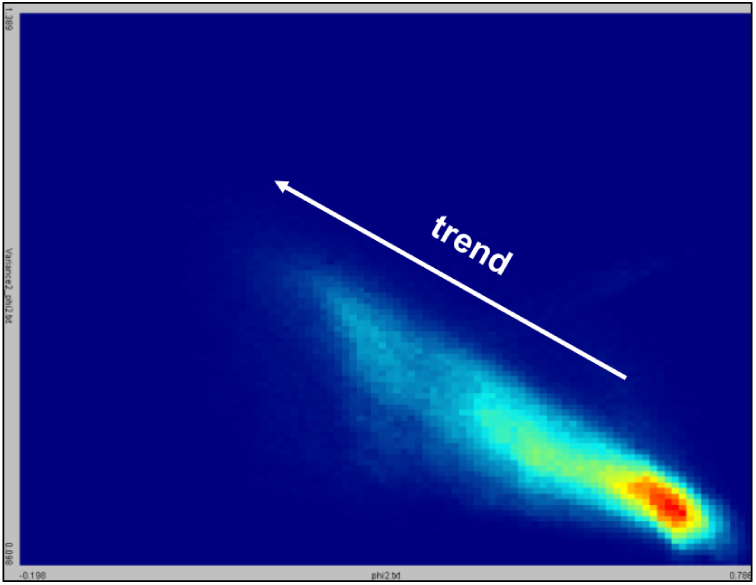
Two phenotype parameters visualized in a 2D histogram. The 2D histogram is a heat map where each cell is colored based on the number of data points that fall into the corresponding bin. An animation is built showing how the relationship between the two phenotype parameters changes over time.

**Fig. 11.**
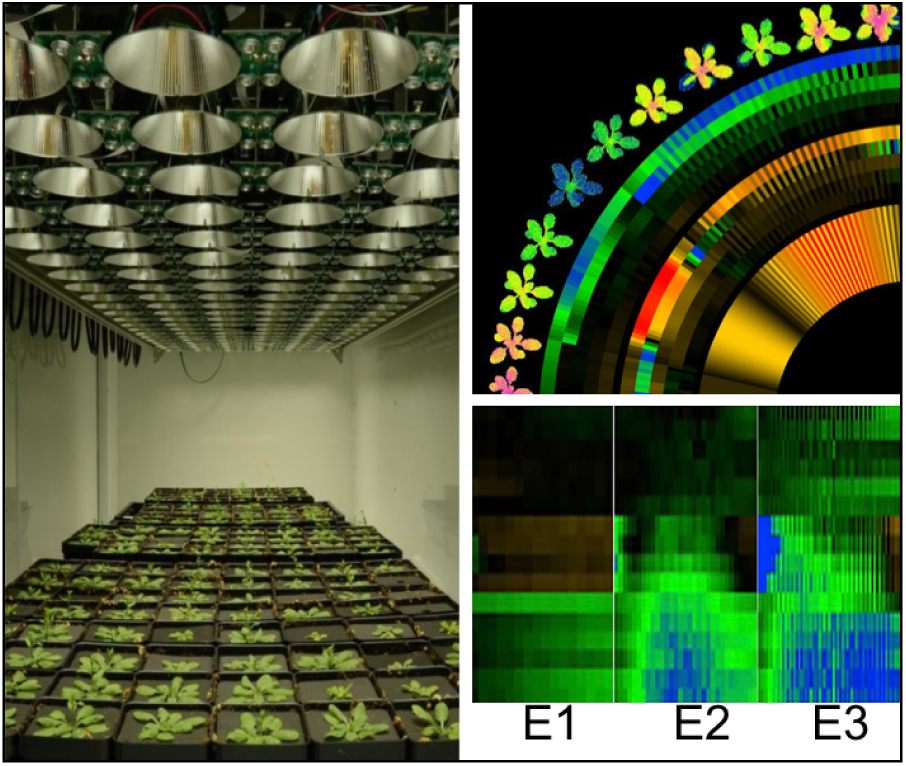
A use case of OLIVER for large-scale plant photosynthetic performance phenotyping [7]. (Left panel) the plant photosynthetic performance of *Arabidopsis thaliana* mutant lines was monitored in a DEPI chamber under three kinds of ramped environmental perturbations (i.e. *E*1, *E*2, and *E*3). (Bottom-right panel) a quick visualization using OLIVER indicates that the emergent phenotypes in the Arabidopsis mutants are highly transient and heterogeneous, depending in complex ways on both environmental conditions and plant developmental age. (Top-right panel) a circular heat map with falsecolor plant images generated using OLIVER, indicating that such emergent phenotypes appear in different leaves under different conditions.

#### 3) Animation

Gaining insights from phenomics data typically requires visualizing and operating upon multiple heat maps. OLIVER can generate a 2D-histogram (see Figure 10) to visualize the temporal correlations between two heat maps using animation. The 2D-histogram itself is a heat map where each cell is a bin, where the coordinates of a bin represents the values in the first and second heat maps, and the color of a bin is proportional to the number of phenotype values in the bin. The animation of temporal correlation can be adjusted in real-time and the results can be exported using the graphics interchange format (GIF) showing how the relationship between two phenotypes changes over time. The entire workspace can also be captured in animation, in which a sliding-window selection of heat map rows is gradually shifted between frames. Any related frames visible in the workspace (e.g. other heat maps, plots, and histograms) are automatically updated for each frame and are included in the animation.

## IV. RESULTS AND DISCUSSION

Use cases in plant phenomics were collected to demonstrate the functionality of OLIVER. In this article, we report a typical use case with large-scale phenomics data and present more use cases in the OLIVER website.

Figure 11 shows a use case of OLIVER for large-scale plant photosynthetic performance phenotyping [7]. The left panel shows that the plant photosynthetic performance of more than 100 *Arabidopsis thaliana* mutant lines was monitored in a DEPI chamber under ramped light intensity perturbations. A quick visualization of the phenomics data using OLIVER (shown in the bottom-right panel) indicates that the emergent phenotypes in the Arabidopsis mutant lines are highly transient and heterogeneous, depending in complex ways on both environmental conditions and plant developmental age. Furthermore, the top-right panel shows a circular heat map with false-color plant images generated using OLIVER, indicating that such emergent phenotypes appear in different leaves under different conditions. These emergent phenotypes appear to be caused by a range of phenomena, suggesting that such previously unseen processes are critical for plant responses to dynamic environments [7].

## V. CONCLUSION

High-throughput phenotyping has revolutionized biological sciences. The power of new phenotyping techniques lies in the fact that they enable the large-scale high-resolution observation of phenotypes, which is key to understanding the molecular mechanisms underlying gene functions in phenotype-related pathways. As a consequence, the quantity of phenomics data has been expanding exponentially in the last a few years.

Analyzing longitudinal phenomics data requires approaches and tools that are distinct from those used to analyze more static genomic data, since phenotyping platforms can aggregate results from a wide range of different sensors, each with its own set of constraints, time resolution and sensitivities. OLIVER is a visual phenomics data analysis tool that integrates data visualization, statistical tests, and real-time data operation, providing researchers an interactive tool, aimed at establishing a comprehensive integration of thoughts, knowledge and data emerge from the interactive activities.

Note the two key differences between OLIVER and the existing heat map tools are that 1) any user adjustment will trigger real-time display updates for any affected elements in the workspace, and 2) multiple heat maps can be synchronously adjusted, operated and connected. These allows scientists to instantly compare multiple phenomics data sets resulting from the same large-scale experiments, or correlate them with genomics information in public databases, in an interactive visualization environment through heat map views. By creating unique images and diagrams to effectively convey messages to scientists, OLIVER will assist them to quickly integrate, visualize, and compare large phenomics data, thus enabling new biological discoveries.

## VI. SOFTWARE AND DATA AVAILABILITY

The OLIVER program, sample data and the user’s manual are available under the the GNU General Public License at the OLIVER website https://caapp-msu.bitbucket.io/projects/oliver/. The latest version of Java should be installed before running OLIVER.

